# Thalamocortical contribution to cognitive task activity

**DOI:** 10.1101/2022.06.28.497905

**Authors:** Evan Sorenson, James M. Shine, Michael W. Cole, Kai Hwang

**Author notes:** **Corresponding Author:** Kai Hwang, Ph.D., Department of Psychological and Brain Sciences, The University of Iowa, G60 PBSB, 340 Iowa Ave, Iowa City, IA 52245. The authors declare no competing financial interests.

## Abstract

Thalamocortical interaction is a ubiquitous functional motif in the mammalian brain. Previously (Hwang et al., 2021), we reported that lesions to network hubs in the human thalamus are associated with multi-domain behavioral impairments in language, memory, and executive functions. Here we show how task-evoked thalamic activity and thalamocortical interactions are organized to support these broad cognitive abilities. To address this question, we analyzed functional MRI data from human subjects that performed 127 tasks encompassing a broad range of cognitive representations. We first investigated the spatial organization of task-evoked activity and found that multi-task thalamic activity converged onto a low-dimensional structure, through which a basis set of activity patterns are evoked to support processing needs of each task. Specifically, the anterior, medial, and posterior-medial thalamus exhibit hub-like activity profiles that are suggestive of broad functional participation. These thalamic task hubs overlapped with network hubs interlinking cortical systems. To further determine the cognitive relevance of thalamocortical interactions, we built a data-driven thalamocortical interaction model to test whether thalamocortical functional connectivity transformed thalamic activity to cortical task activity. The thalamocortical model predicted task-specific cortical activity patterns, and outperformed comparison models built on cortical, hippocampal, and striatal regions. Simulated lesions to low-dimensional, multi-task thalamic hub regions impaired task activity prediction. This simulation result was further supported by profiles of neuropsychological impairments in human patients with focal thalamic lesions. In summary, our results suggest a general organizational principle of how thalamocortical interactions support cognitive task activity.

**Impact Statement:** Human thalamic activity transformed via thalamocortical functional connectivity to support task representations across functional domains.

## Introduction

Distributed neural activity supports a broad range of perceptual, motor, affective, and cognitive functions. Discovering how brain systems implement this broad behavioral repertoire is critical for elucidating the neural basis of human cognition. Past studies have revealed two organizational principles—low-dimensional architecture and multi-task hubs—that connect distributed neural activity with task performance across functional domains.

The application of dimensionality reduction techniques on whole-brain imaging data has revealed a relatively low-dimensional organization of distributed neural activity. Specifically, the number of variables required to explain a large amount of variance in distributed neural activity is far lower than the total number of variables in the data (MacDowell and Buschman, 2020; Nakai and Nishimoto, 2020; Shine et al., 2019a). The spatiotemporal patterns of these variables are commonly referred to as intrinsic networks in task-free conditions, or manifolds, motifs, and latent components in task contexts. This low-dimensional organization may reflect an elementary set of information processes implemented by distributed brain systems (Cunningham and Yu, 2014; Yeo et al., 2016), in which different tasks selectively engaged selective latent activity patterns depending on the specific processing requirements of individual tasks.

The anatomical overlap between spatiotemporal components predicts that some brain regions broadly participate in multiple tasks. In support of this prediction, studies have found that associative regions in frontal and parietal cortices are involved in executing a wide array of tasks (Cole et al., 2013; Duncan, 2010). These task-flexible regions, also commonly referred to as brain hubs (Gratton et al., 2018; van den Heuvel and Sporns, 2013), have diverging connectivity with multiple brain systems, and are thought to perform integrative functions that allow perceptual inputs to interact with contextual task representations for adaptive task control (Bertolero et al., 2018, 2015; Ito et al., 2022; Nee, 2021). The behavioral significance of brain hubs is affirmed by lesion studies demonstrating that lesions to hub regions are associated with task impairments across multiple functional domains (Hwang et al., 2021; Reber et al., 2021; Warren et al., 2014).

However, a prevailing assumption is that diverse human behavior depends on the organization of cortical activity, and the contribution from subcortical regions, particularly the thalamus, is not well understood. Our previous studies demonstrated that the human thalamus contains a complete representation of intrinsic cortical functional networks, and additionally, exhibits a hub-like connectivity profile interlinking multiple cortical systems (Hwang et al., 2021, 2017). Furthermore, lesions to the anterior and the medial thalamus in human patients are associated with behavioral impairments across functional domains (Hwang et al., 2020a, 2021). Given that every cortical region receives projections from multiple thalamic nuclei and that the thalamus mediates striatal and cerebellar influences on cortical processes (Jones, 2001; Sherman, 2007; Shine, 2020), thalamocortical interaction is ideally suited to shape cortex-wide activity patterns that instantiate cognitive representations. Whether thalamic task activity exhibits similar low-dimensional organization and how thalamocortical interactions contributes to cognitive task activity remains unclear.

The goal of the current study was to determine how thalamic task-evoked activity, low-dimensional organization, and thalamic hub architecture are organized to support cognitive functions across diverse functional domains. To this end, we analyzed functional magnetic resonance imaging (fMRI) data where human subjects performed a rich battery of tasks designed to elicit neural activity for a wide range of cognitive functions. Our study addressed the following three specific questions. First, does task-evoked thalamic activity exhibit a low-dimensional organization? Second, are there task-flexible thalamic hub regions that exhibit task-evoked responses across diverse task domains? Third, can thalamic task-evoked activity transformed via thalamocortical functional connectivity (FC) predict cortical task-evoked activity patterns? Answering these three questions would reveal a general principle of how thalamocortical interactions contribute to higher-order, multi-domain cognitive task activity for flexible human behavior.

## Results

To determine the organization of task-evoked thalamic activity and thalamocortical interactions, we analyzed two publicly available fMRI datasets, both of which utilized rich batteries of tasks designed to elicit a wide range of processes encompassing perceptual, affective, memory, social, motor, language, and cognitive domains (Figure 1A). In the first dataset, the multi-domain task battery (MDTB) dataset, 21 subjects performed 25 behavioral tasks (King et al., 2019). In the second dataset, the Nakai and Nishimoto (N&N) dataset, 6 subjects performed 103 tasks (Nakai and Nishimoto, 2020).

**Figure 1.**
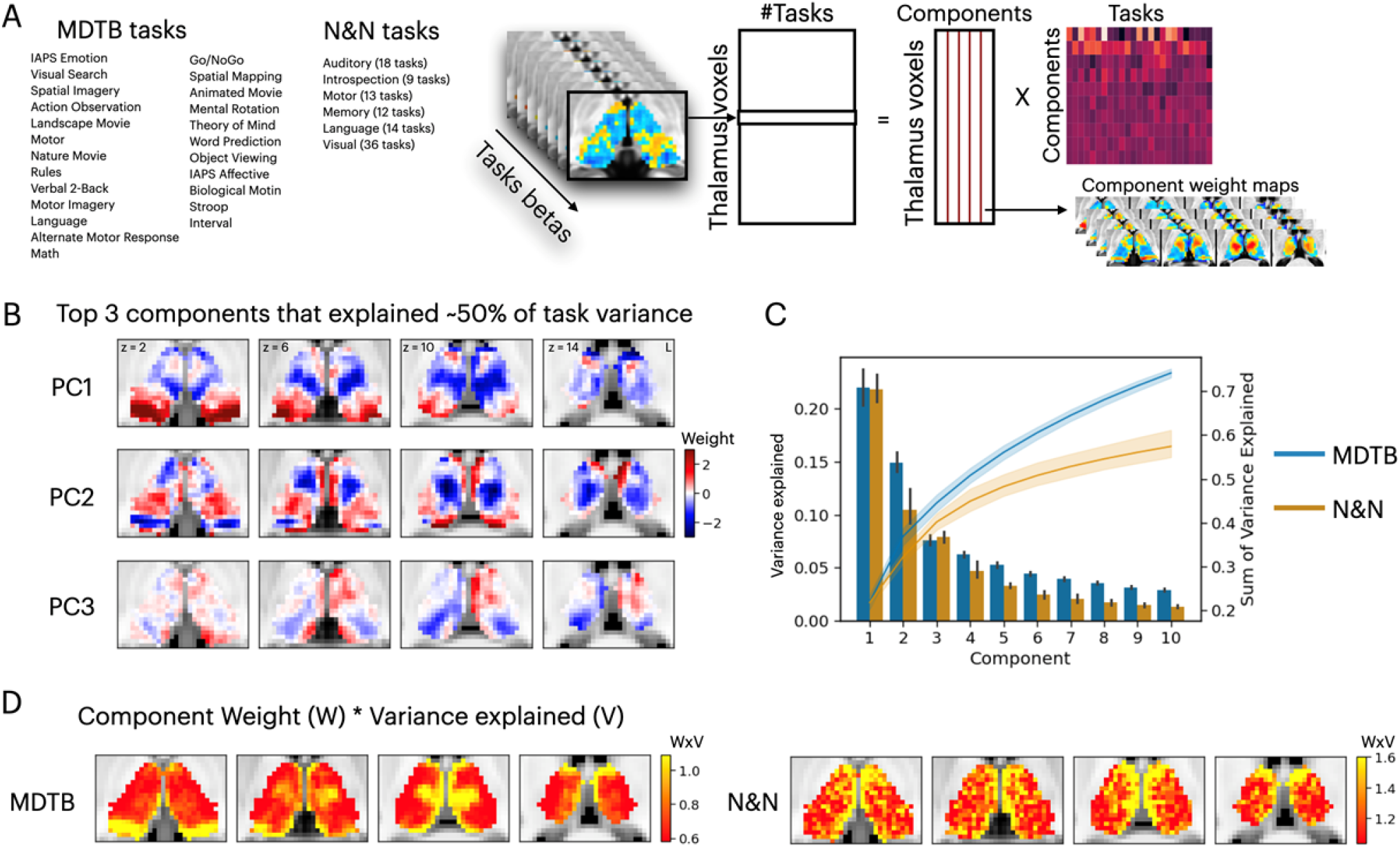
Low-dimensional organization of thalamic task-evoked response that supports multi-task performance. (A) We decomposed the high-dimensional multi-task evoked activity matrix into low-dimensional spatial components in the human thalamus and a task-wide loading matrix. For a list of all tasks see Figure 1-Supplementary File 1 (B) Spatial topography of the top three components from the MDTB dataset that explained about 50% of the variance in the group averaged task activity matrix. For top components for the N&N dataset see Figure 1-Figure Supplement 1. For the loadings between tasks and components, see Figure 1-Figure Supplement 2. (C) Results from applying PCA to single subjects. For both the MDTB and N&N datasets, for individual subjects up to 60% of the variance across multiple tasks can be explained by the top 10 components. Error bars and shaded areas indicate 95% bootstrapped confidence intervals.

### Low dimensional organization of thalamic task-evoked activity

We first sought to determine the low-dimensional organization of task-evoked responses in the human thalamus. We used a general linear modeling approach to characterize the task-evoked blood-oxygen-level-dependent (BOLD) activity patterns and estimated the magnitude of BOLD evoked responses for every task (“task betas”). Task betas were extracted for every thalamic voxel and every task and compiled into a voxel-by-task activity matrix for each dataset. The cross-subject averaged matrix was subjected to a principal component analysis (PCA) to decompose multi-task evoked activity patterns into a linear summation of a voxel-by-component weight matrix multiplied by a component-by-task loading matrix (Figure 1A). The voxel-by-component weight matrix can be conceptualized as sets of basis patterns of thalamic activity components engaged by different tasks. Similar to previous studies that focused on cortical activity patterns (Nakai and Nishimoto, 2020; Shine et al., 2019a), this analysis revealed a low-dimensional organization, in which the top thalamic activity components can explain up to 60% of the variance across tasks, and each task is associated with a weighted sum of these components (Figure 1B, Figure 1 – Figure Supplements 1-2). Repeating the PCA for each subject (without averaging across subjects) showed that this low-dimensional organization is also observed in individual subjects, where variances across task-evoked activity patterns in the human thalamus can be explained by a low number of components relative to the larger number of cognitive tasks administered (Figure 1C).

### Hub organization of thalamic task-evoked activity

Examining the spatial patterns of thalamic components (Figure 1B) revealed that several thalamic subregions showed spatial overlap across multiple components, which suggests that these thalamic subregions may flexibly support multiple tasks. We reasoned that if a thalamic voxel is broadly participating in multiple functional domains, it will express a strong weight in components that explained more variances across tasks. Therefore, to further map task-flexible regions in the human thalamus, we calculated a task hub metric by summing each voxel’s absolute component weight multiplied by the percentage of variance explained by each component. Results showed that the anterior, medial, medio-posterior, and dorsal thalamus exhibited strong task hub properties (Figure 2A); we had also previously identified these same thalamic subregions as network hubs (Hwang et al., 2017).

**Figure 2.**
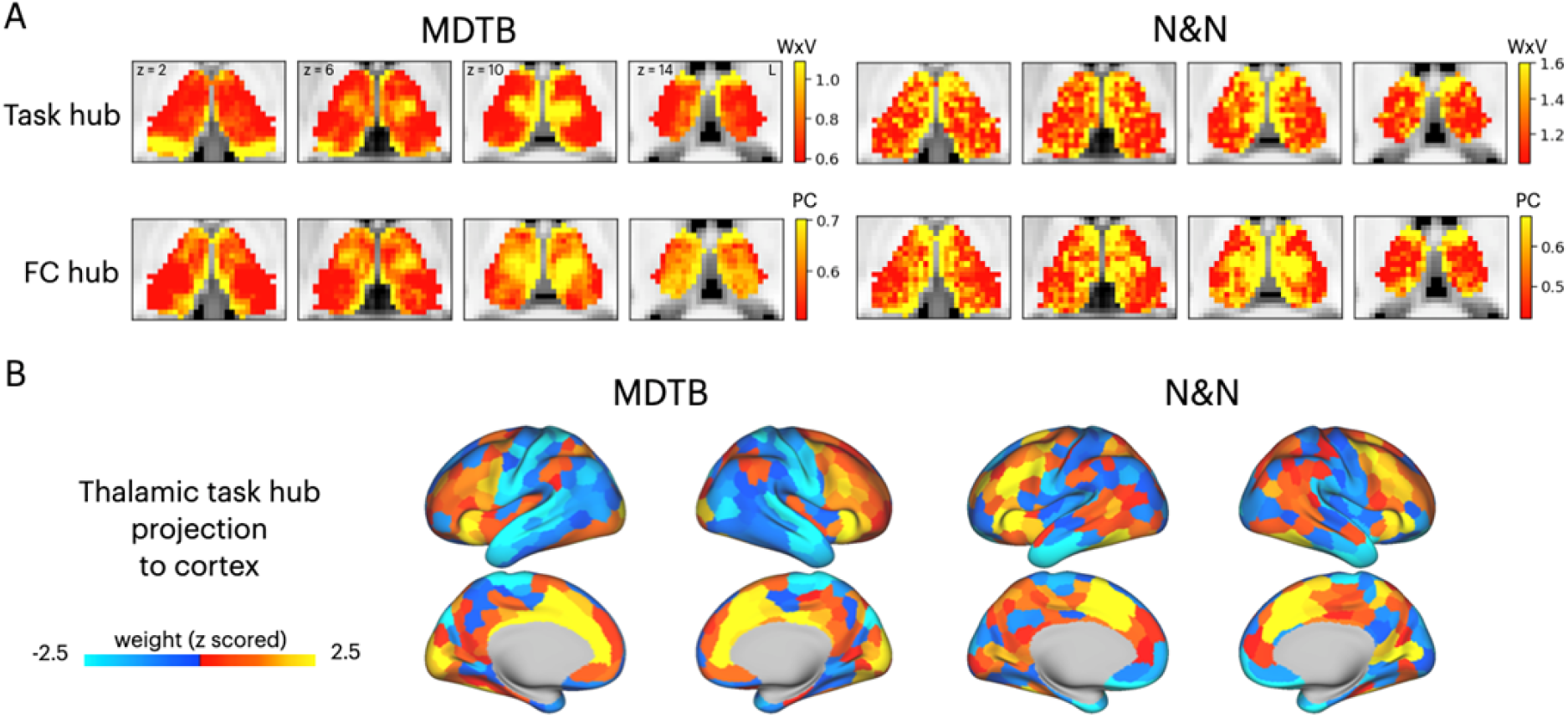
Hub regions in the thalamus. (A) Task hubs and FC hubs in the thalamus. WxV = component weight x variances explained by each component. PC = participation coefficient. (B). Projecting task hub metrics onto the cortex via thalamocortical FC.

Network hubs in the human thalamus exhibit diverse functional connectivity patterns interlinking multiple cortical systems (Greene et al., 2020; Hwang et al., 2017). To determine whether the tasks hubs we identified correspond to network hub regions previously identified in the thalamus, we compared the spatial similarity of task hubs with functional connectivity hubs (FC hubs). We calculated a network hub metric, the participation coefficient (Gratton et al., 2012; Guimerà and Nunes Amaral, 2005), for every thalamic voxel using timeseries after removing task-evoked variances from the preprocessed fMRI data. The task hubs showed significant spatial correspondence with FC hubs (Figure 2A; spatial correlation: MDTB mean = 0.17, SD = 0.083, p < 0.001; N&N mean = 0.17, SD = 0.05, p < 0.001). One discrepancy we identified was the posterior thalamus, which showed strong task hub but not network hub property for the MDTB dataset. We then projected the task hubs estimates onto the cortex by calculating the dot product between each thalamic voxel’s task hub estimate and its thalamocortical FC matrix. We found that tasks hubs were most strongly coupled with associate regions in the frontal and parietal cortex, for example the lateral frontal, the insula, the dorsal medial prefrontal, and the intraparietal cortices (Figure 2B), regions previously identified as cortical network hubs (Bertolero et al., 2018, 2015).

### Thalamocortical interactions transform cognitive task activity

To further test whether thalamic task-evoked activity can support diverse cognitive functions putatively implemented by distributed cortical activity, we adapted the activity flow mapping procedure (Cole et al., 2016; Ito et al., 2017) to assess whether thalamocortical interactions can plausibly transform thalamic task-evoked responses to cortical task activity (Figure 3A). Briefly, for every thalamic voxel, we calculated the dot product between the thalamic evoked response pattern of each task and the thalamocortical FC matrix. This calculation yielded a predicted cortex-wide activity pattern for every task, which can be compared to the observed cortical activity pattern using Pearson correlation. One null model randomly shuffled thalamic evoked responses (the “null model”) while the other one set all thalamic evoked responses to a uniform value across all voxels (the “uniformed evoked model”). Both null models assumed no spatial structure in thalamic task-evoked responses, and the uniformed evoked model further assumed that cortical activity patterns are determined only by the summated inputs from thalamocortical FC patterns. We found that across tasks, our thalamocortical activity-flow model outperformed the null models (Figure 3B; MDTB thalamocortical model: mean pattern correlation across tasks = 0.39, SD = 0.1; MDTB null model: mean r = 0.09, SD = 0.053; MDTB uniformed evoked model: mean r = -0.002, SD = 0.11; MDTB thalamocortical model vs null model: t(24) = 15.27, p < 0.001; MDTB thalamocortical model vs uniformed evoked model: t(24) = 14.13, p < 0.001; N&N thalamocortical model: mean pattern correlation across tasks = 0.23, SD = 0.064; N&N null model: mean r = 0.073, SD = 0.044; N&N uniformed evoked model: mean r = -0.021, SD = 0.22; N&N thalamocortical model vs null model: t(102) = 28.4, p < 0.001; N&N thalamocortical model vs uniformed evoked model: t(102) = 17.98, p < 0.001). These results indicate that thalamocortical interactions carry task information.

**Figure 3.**
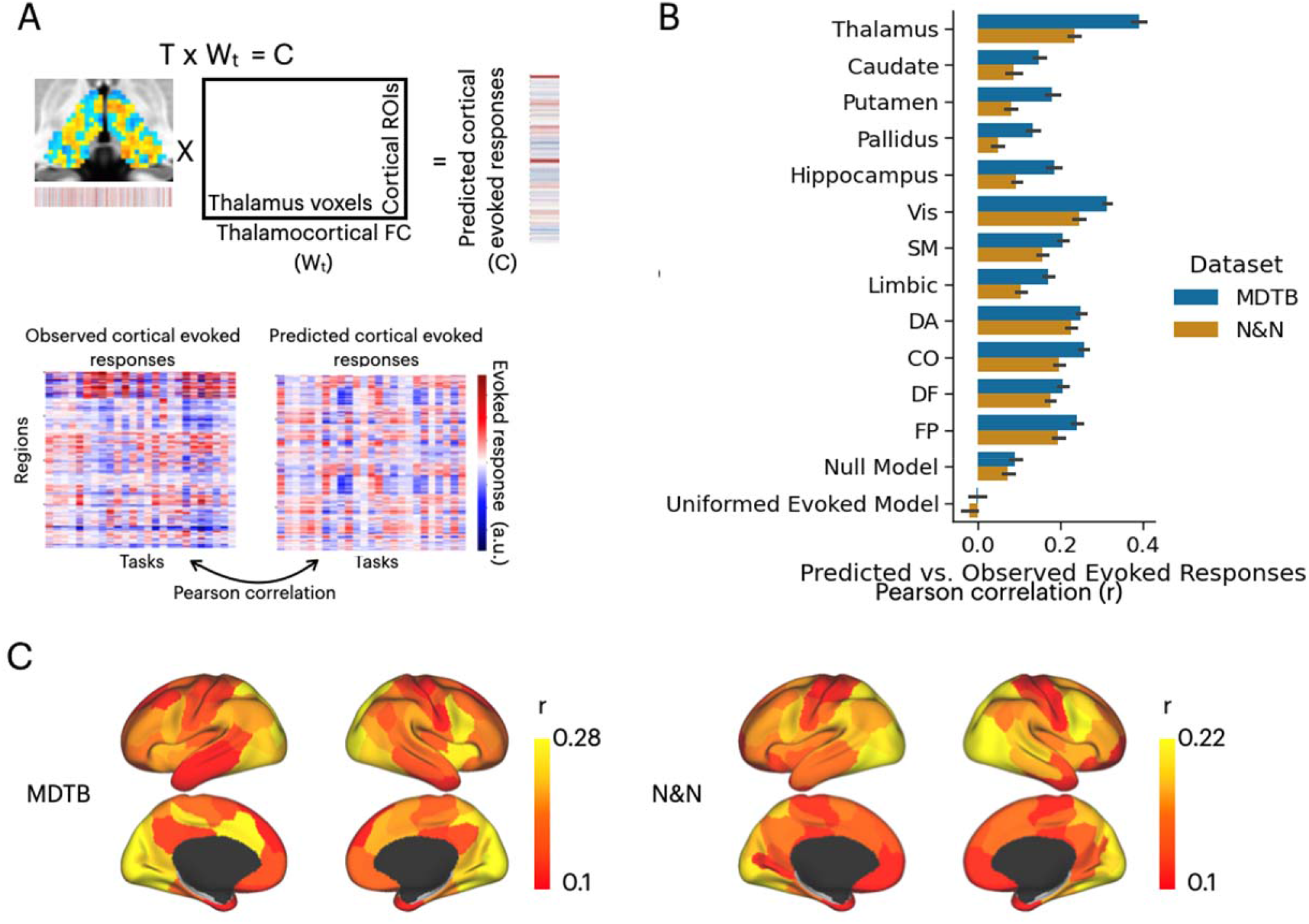
Thalamus multi-task activity and its influence on multi-task cognitive activity in the cortex. (A) Model for testing whether patterns of cortical task-evoked responses can be predicted by thalamic task-evoked activity transferred through thalamocortical FC. (B) Compared to other brain structures, thalamic task-evoked activity has the strongest predictive power of multi-task cortical activity patterns. Cortical regions divided by 7 cortical functional networks: Vis = visual, SM = somatomotor, DA = dorsal attention, CO = cingulo-opercular, DF = default mode, FP = frontal parietal; for a detailed list of prediction results for every ROI and every task see Figure 3-Supplemental data 1. Null model: randomly permute the evoked pattern in the thalamus. Uniformed model: setting thalamic evoked responses to uniform across all voxels. Error bars indicate 95% bootstrapped confidence intervals. (C) In addition to the thalamus, the visual cortex, the insula, the inferior frontal cortex, the intraparietal sulcus, the medial frontal cortex, and the precentral sulcus also showed strong activity predictions.

We then sought to compare our thalamocortical activity flow model to other cortical and subcortical regions. We constructed a series of comparison models by applying the same activity flow mapping procedure to 100 cortical regions of interests (ROIs) that had similar size as the thalamus (Schaefer et al., 2018), as well as to other subcortical regions including the hippocampus, the caudate, the putamen, and the globus pallidus (Figure 3B). We found that several cortical regions also showed strong prediction performance, including the insula, the inferior frontal cortex, the dorsal medial frontal cortex, the intraparietal sulcus, and the visual cortex (Figure 3C; For a detailed list of prediction results for every ROI and every task, see Figure 3-Supplement File-1). However, the thalamocortical model outperformed all comparison models except cortical ROIs from the visual cortex from the N&N data. These results indicate that thalamocortical interaction is one of the strongest predictors of cortical task activity.

### Representation geometry predicted from thalamocortical interactions

In addition to predicting activity patterns for individual tasks, we also tested whether thalamocortical activity-flow can predict the dissimilarity among task activity patterns, a measure also known as representational geometry (Kriegeskorte and Kievit, 2013). Specifically, representation geometry seeks to capture the topography of task representations by characterizing the multivariate distance among activity patterns for each task. We calculated the representational dissimilarity matrix (RDM) using Spearman correlation from both the predicted and observed cortical activity patterns (Figure 4A). Predicted and observed RDMs were then compared using Kandal’s Tau (Nili et al., 2014). We found that across subjects, the thalamocortical activity-flow model performed significantly better than the two null models (MDTB thalamocortical model: mean tau = 0.34, SD = 0.11; MDTB null model: mean r = 0.17, SD = 0.077; MDTB uniformed evoked model: mean r = 0.012, SD = 0.037; MDTB thalamocortical model vs null model: t(20) = 8.35, p < 0.001; MDTB thalamocortical model vs uniformed evoked model: t(20) = 13.82, p < 0.001; N&N thalamocortical model: mean tau = 0.18, SD = 0.064; N&N null model: mean r = 0.12, SD = 0.054, N&N uniformed evoked model: mean r = - 0.0047, SD = 0.27; N&N thalamocortical model vs null model: t(5) = 7.02, p < 0.001; N&N thalamocortical model vs uniformed evoked model: t(5) = 5.84, p = 0.002), and outperformed most comparison models except the visual cortex and cortical regions in the DA network for the N&N dataset (Figure 4B; For a detailed list of prediction results from all ROIs and tasks see Figure 4-Supplementary File-1). We note that the predicted RDM suggests the thalamocortical activity flow model showed differences in activity patterns between tasks, as the predicted distances varied. In addition to the thalamus, comparison models constructed with activity patterns from the insula, the dorsal medial prefrontal cortex, the precentral sulcus, the inferior frontal cortex, the intraparietal sulcus, and the visual cortex also showed strong prediction of representational geometry, but weaker than the thalamus (Figure 4B-C). These results indicate that thalamocortical interaction provides one of the strongest contributions to cortical representational geometry across multiple functional domains.

**Figure 4.**
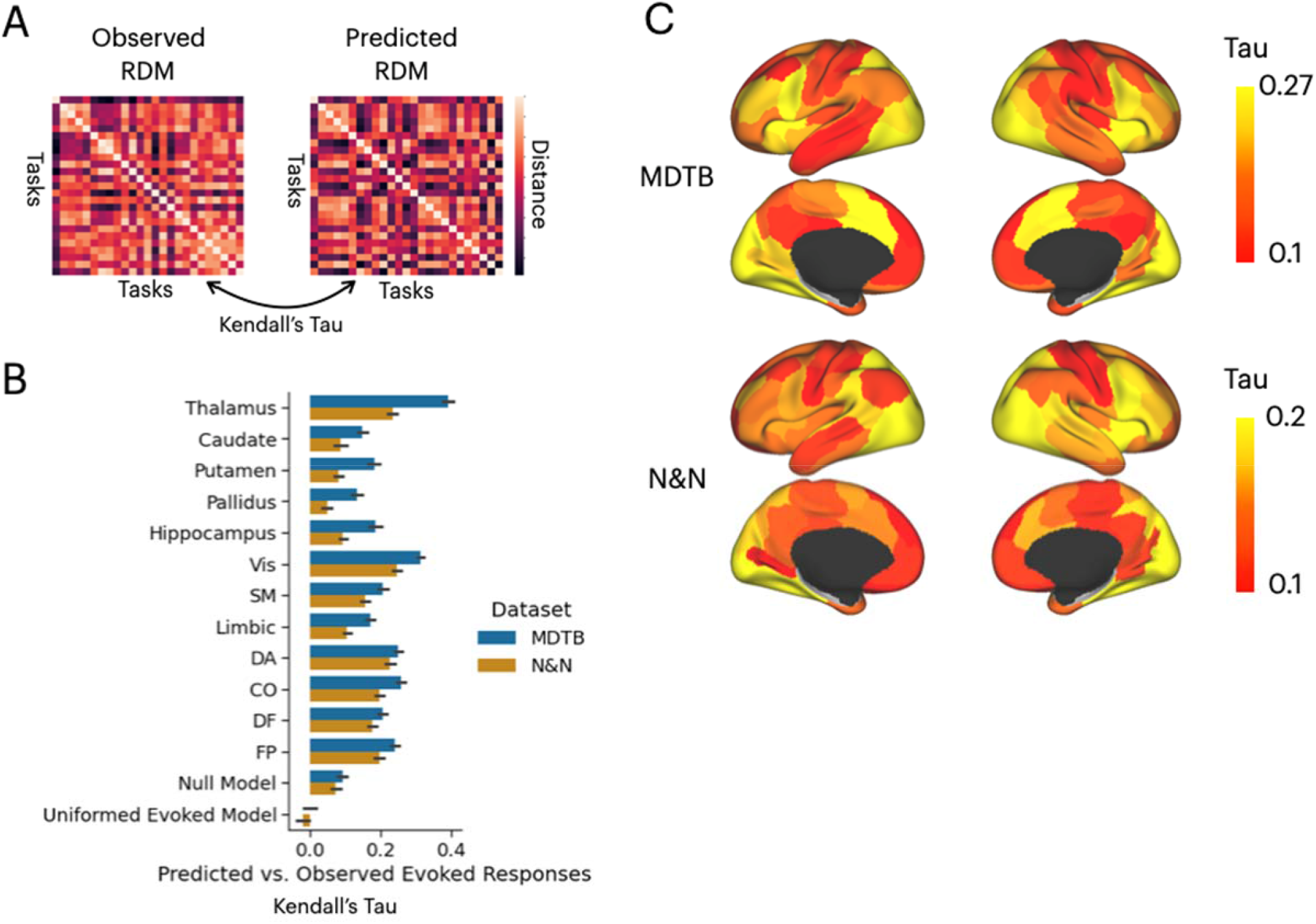
Thalamus multi-task activity and its influence on multi-task representational geometry in the cortex (A) Model for testing whether multi-task representational geometry constructed from observed cortical responses can be predicted by thalamocortical activity flow (Figure 3) (B) Compared to other brain structures, thalamic task-evoked activity showed stronger predictive power of multi-task representational geometry than most comparison models. Cortical regions divided by 7 cortical functional networks: Vis = visual, SM = somatomotor, DA = dorsal attention, CO = cingulo-opercular, DF = default mode, FP = frontal parietal. For a detailed list of prediction results for every ROI and every task see Figure 4-Supplemental data 1. Null model: randomly permute the evoked pattern in the thalamus. Uniformed model: setting thalamic evoked responses to uniform across all voxels. Error bars indicate 95% bootstrapped confidence intervals. (C) In addition to the thalamus, the visual cortex, the insula, the inferior frontal cortex, the medial frontal cortex, the intraparietal sulcus, and the precentral sulcus also showed strong predictions.

### Lesions to task hubs in the thalamus are associated with impaired prediction of task reorientations and representational geometry

The results presented in figures 3 and 4 demonstrated that thalamic task-evoked responses can be transformed to cortical activity patterns to putatively support cognitive representations via thalamocortical interactions. However, it is unclear to what degree task hubs in the thalamus (Figure 3A) contribute to these results. One possibility is that all thalamic voxels contribute equally to cortical task activity patterns. Another possibility is that, since thalamic task hubs are broadly engaged by multiple tasks and exhibited diverging connectivity patterns with multiple cortical systems, thalamic task hubs have a stronger contribution to predicting cortical task activity patterns. To evaluate this prediction, we performed virtual lesion simulations. Specifically, we systematically removed 20% of thalamic voxels based on their percentile rank of task hub estimates and calculated the percentage of reduction in prediction accuracy from the activity-flow analysis. For both datasets, we found that removing voxels with strong task hub estimates decreased the prediction accuracy in cortical task-evoked activity patterns (Figure 5A) and representational geometry (Figure 5C). We fitted a regression model to test the relationship between reduction in prediction and percentile rank of task hub metrics removed, and found significant negative associations (MDTB activity flow prediction: b = -0.43, SE = 0.019, t = -22.63, p < 0.001; N&N activity flow prediction: b = -0.1, SE = 0.028, t = -3.79, p < 0.001; MDTB RDM prediction: b = -0.61, SE = 0.029, t = -21.3, p < 0.001; N&N RDM prediction: b = - 0.6454, SE = 0.066, t = -9.72, p < 0.001). These results indicate that virtual lesioning of thalamic task hubs had a stronger impact on reducing the model’s ability to predict cortical activity patterns. Furthermore, thalamic voxels with the strongest impact on prediction performance were spatially located in the anterior, medial, mediodorsal, and medial posterior thalamus that we previously identified as task hubs (Figure 5B and Figure 5D).

**Figure 5.**
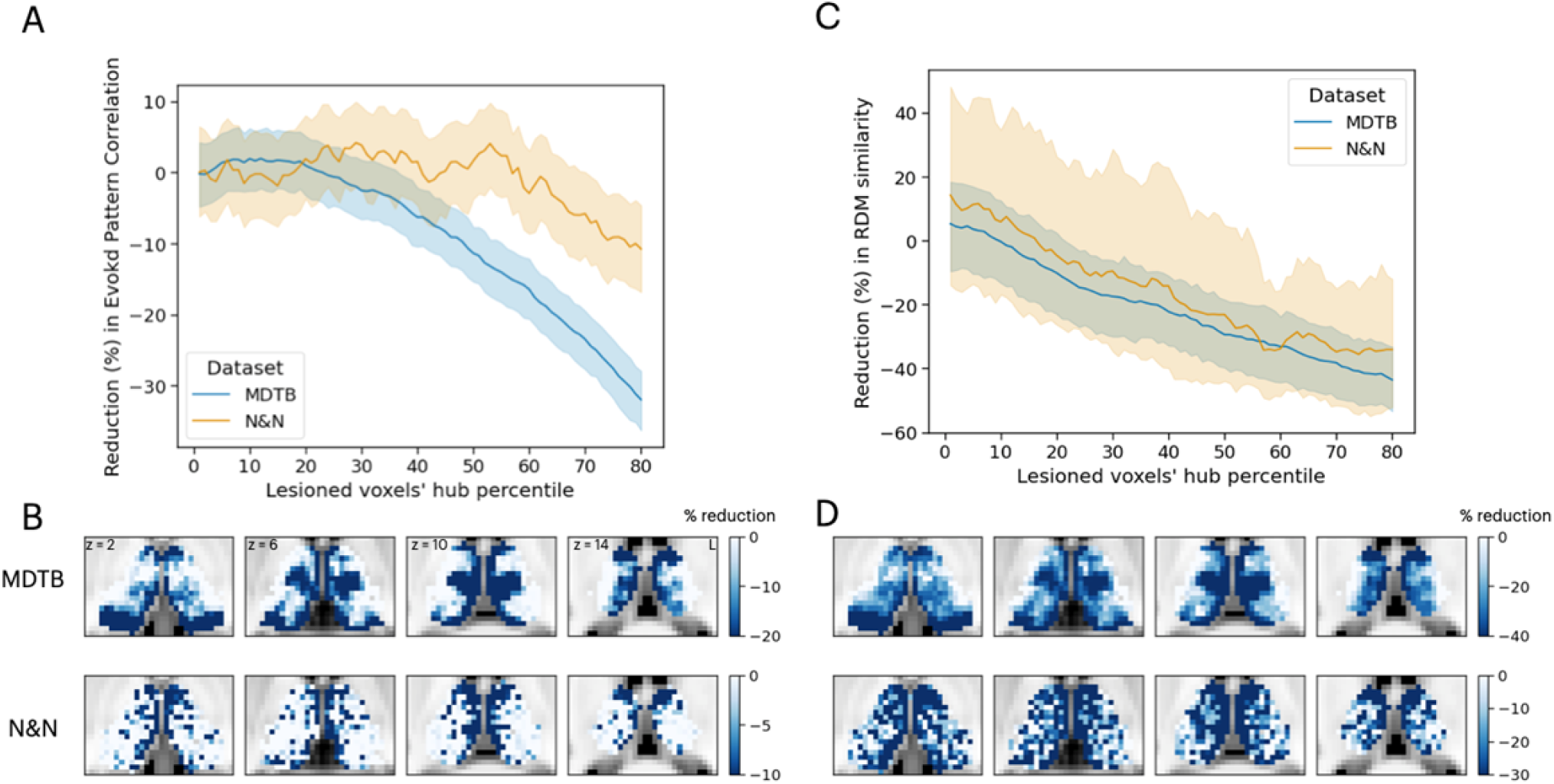
Simulating the thalamic lesion’s effect on multi-task evoked pattern similarity and representational geometry. (A) Artificial lesion of 20% of the thalamus voxels based on their percentile rank of task hub property: examination of the impact on task activity prediction. (B) Subregions that showed greater reduction in prediction accuracy were primarily located in anterior, medial, and posterior thalamus. (C) Artificial lesion of 20% of the thalamus voxels based on their percentile rank of task hub property: examination of the impact on representation geometry prediction. (D) Subregions that showed greater reduction in prediction accuracy of representation geometry showed spatial correspondence with task hubs in the thalamus. Shaded error indicates 95% bootstrapped confidence intervals.

We compared these simulation results with neuropsychological impairments found in 20 human patients with focal thalamic lesions. Neuropsychological profiles of these patients were described in detail in our previous publication (Hwang et al., 2021). Briefly, patients performed a battery of neuropsychological tests to assess executive, language, memory, learning, visuospatial, and construction functions (Lezak et al., 2012). Test performance was then compared to published, standardized norms and converted to z-scores quantify the severity of impairment (Figure 6A). Patients were grouped into two groups, those that showed impairment (z-score < -1.695, worse than 95 percentile of the normative population) in single or fewer domains (the SM group) versus those showed impairment across multiple domains (the MM group; Figure 6B). There were no statistically significant differences in the lesion volumes between these two groups of patients (MM group: mean = 1559 mm^3^, SD = 1370 mm^3^; SM group: mean = 1073 mm^3^, SD = 925 mm^3^; group difference in lesion size, t(18) = 0.87 p = 0.39).

**Figure 6.**
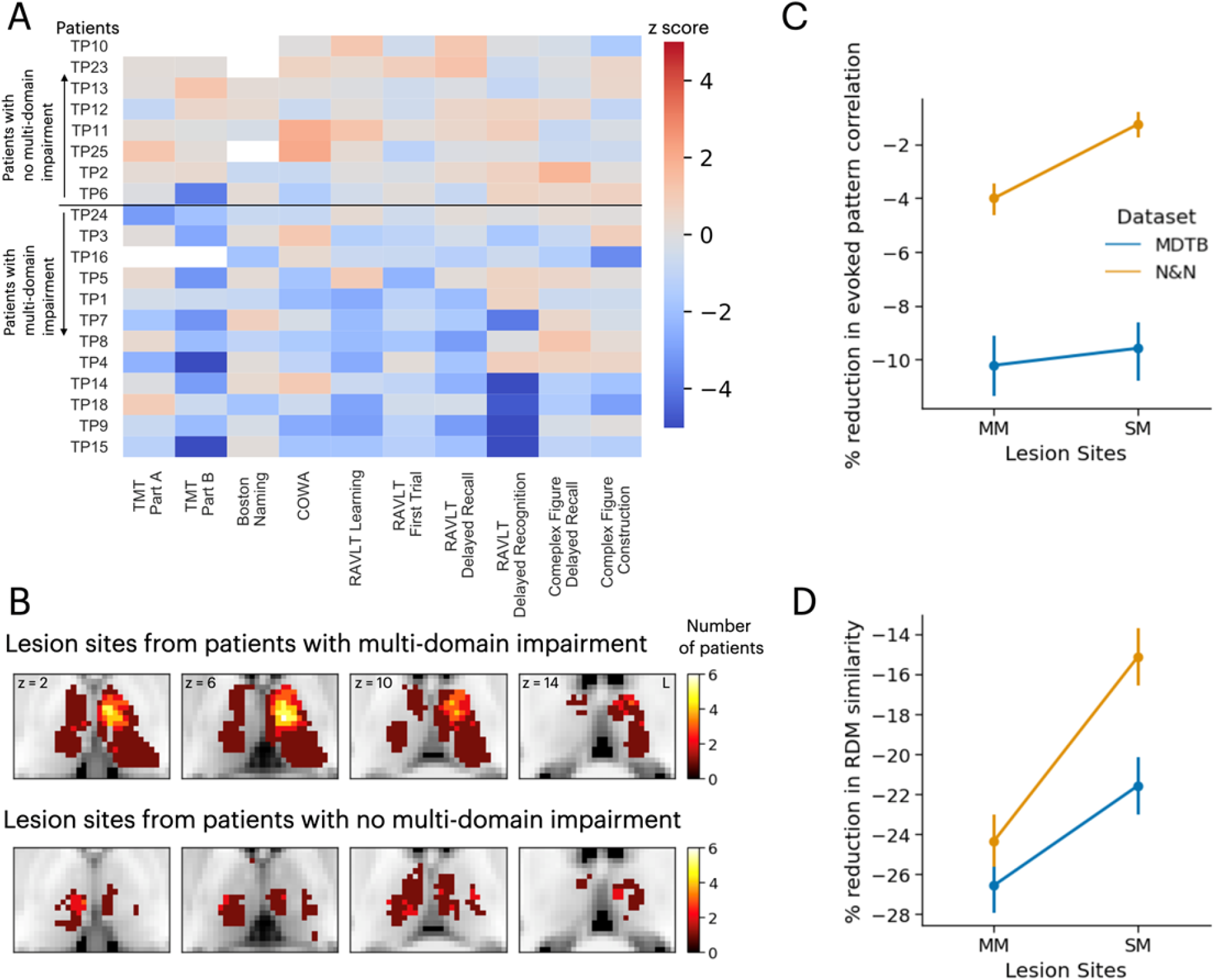
Neuropsychological evaluations from 20 patients with focal thalamic lesions. (A) Twelve patients exhibited multi-domain (MM) impairment (negative z-scores) across multiple neuropsychological assessments. Eight patients exhibited no multi-domain (SM) impairment. (B) Lesion sites from patients with and without multi-domain impairment. (C-D) Mean and 95% bootstrapped confidence interval of the reduction activity prediction after virtual lesions, plotted separately for virtual lesion sites that overlapped with MM or SM lesions. Lesion sites associated with multi-domain impairment also associated with a larger reduction in multi-task pattern similarity after simulated lesions (C) and in representational geometry (D). Figure 6A reproduced from Hwang et al., 2021 with permission.

Lesion sites from each group were compared to the effects of the simulated lesions presented in Figure 6B-D. Specifically, we compared the percentages of reduction in voxels from these two groups of lesions. We found that simulated lesions to voxels that overlapped with lesion sites in the MM patient group showed larger reductions in task activity predictions for the N&N dataset but not the MDTB dataset (Figure 6C; MDTB: Komogorov-Smirnov Test D = 0.048, p = 0.065; N&N: Komogorov-Smirnov Test D = 0.25, p < 0.001). Parallel findings were also observed for representational geometry (Figures 6D; MDTB: Komogorov-Smirnov Test D = 0.17, p < 0.001; N&N: Komogorov-Smirnov Test D = 0.28, p < 0.001). These empirical results from human patients support our simulated lesion analyses, suggesting that lesioning task hubs in the human thalamus are associated with behavioral deficits across functional domains.

## Discussion

It was hypothesized that the thalamus influences cognitive representations beyond sensory and motor domains (Halassa and Sherman, 2019; Wolff and Vann, 2019). Anatomically, every cortical region receives inputs from one or many thalamic subregions, and most thalamic subregions send signals to one or many cortical systems (Jones, 2001; Sherman, 2007). Functionally, resting-state fMRI studies have found a complete representation of intrinsic cortical networks in the human thalamus (Greene et al., 2020; Hwang et al., 2017; Yuan et al., 2016; Zhang et al., 2008). Several of these networks, such as the FP, CO, and DF, have been implicated in multiple cognitive functions (Sadaghiani et al., 2010; Seeley et al., 2007). These observations raised a broader question of if and how the human thalamus contributes to cognitive task activity across functional domains – the central question we sought to address in this study.

For the current study, we investigated how thalamic task-evoked activity and thalamocortical functional interactions are organized. We identified several key organizational characteristics pertaining to these activities. First, similar to cortical task activity, thalamic task-evoked activity exhibited a low-dimensional organization, in which a relatively low number of activity patterns can explain a large amount of the variances observed in multi-task thalamic activity patterns. Second, the anterior, medial, and posterior-medial thalamus were more functionally flexible and participated in multiple tasks, and these “task-hubs” spatially overlapped with FC hubs mapped in our previous studies (Hwang et al., 2021, 2017). Third, thalamocortical functional interactions transform thalamic task-evoked activity to task-specific activity patterns in the cortex. Critically, when compared to other brain structures, this thalamocortical activity flow model performed the best in predicting cortical task activity, further highlighting the capacity of the thalamus in influencing distributed activity patterns that may instantiate cognitive representations. Finally, findings from the thalamocortical activity flow model were further corroborated by simulated lesions and neuropsychological impairments observed in patients with focal thalamic lesions, affirming the behavioral significance of thalamocortical interactions. Collectively, these findings highlight a general and critical role of thalamocortical interactions supporting cognitive functions.

Neural systems support a rich and diverse behavioral repertoire. Findings from both human functional neuroimaging and animal models now point to the low-dimensional structure as a key organizational characteristic that interlinks spatiotemporal neural activity and behavior (Beam et al., 2021; Karolis et al., 2019; MacDowell and Buschman, 2020; Nakai and Nishimoto, 2020; Shine et al., 2019a). For example, a large-scale meta-analysis on more than 10,000 neuroimaging studies found that activity patterns across 83 behavioral tasks can be summarized by 13 latent components (Yeo et al., 2016). This study formalized the notion that task performance depends on sets of cognitive components, and that the neuroanatomical distribution of task-evoked activity has a low-dimensional organization. Behaviorally, similar observations have also been reported in the clinical profiles from human stroke patients (Bisogno et al., 2021), demonstrating that three components can explain 50% of behavioral deficits across neuropsychological batteries. Analogous findings have also been reported for the temporal dynamics of neural activities, in which spatiotemporal patterns associated with memory, social, emotion, and language tasks correspond to the temporal flow within a low-dimensional state space (Shine et al., 2019a). Altogether, these studies suggest that an elementary set of processes can be deployed adaptively to support processing demands across different tasks.

However, most prior studies focused on cortical activity patterns, therefore one contribution of our study was to identify a low-dimensional activity structure in the human thalamus. This observation may refine our understanding of thalamic contributions to human cognition. For example, the thalamus itself likely does not encode high-dimensional, vivid, cognitive representations for each individual task. Instead, thalamocortical circuits may support a few key, common but significant, functions across tasks. Notably, the thalamic task components we observed expressed strongest weights in anterior, medial, and posterior thalamus but not in lateral thalamic regions that overlap with first-order thalamic relays such as the lateral geniculate nucleus and the ventrolateral nucleus. This observation suggests that the thalamus likely supports functions beyond basic sensorimotor relay, some of which were suggested by previous studies, and include selection and gain control (Halassa and Kastner, 2017), adjustment of inter-regional communication (Hwang et al., 2020b; Saalmann et al., 2012), and modulation of cortical excitability (Kosciessa et al., 2021). The small size of the thalamus further suggests a prominent role in modulating the low-dimensional organization of task-evoked cortical activity – for instance, previous studies have found that thalamic activity correlates with the dimensionality and the strength of cross-system coupling between cortical regions (Garrett et al., 2018; Shine et al., 2019b). These thalamocortical processes may be required for behavioral performance across many different tasks, driving the low-dimensional organization we observed.

Our findings further support proposals suggesting that the human thalamus, by virtue of its unique connectivity profile, may be in an ideal position to influence cortex-wide task activity (Garrett et al., 2018). Because every cortical region receives inputs from one or multiple thalamic regions, thalamocortical interactions may be more effective in pushing cortical activity patterns to the desired task state. We speculate that the thalamus may be at the nexus where specific activity patterns in the thalamus can be selectively down- or up-regulated to meet the specific information processing demands of different tasks. This hypothesis predicts that thalamocortical interactions should transfer task-specific information between the thalamus and the cortex, a prediction we tested in our thalamocortical activity flow analysis. Specifically, we constructed an activity flow mapping model to test whether thalamocortical interactions can transfer thalamic task-evoked responses to cortical task activity. We found that our data-driven thalamocortical model can indeed successfully predict task-specific activity patterns and multi-task representational geometry observed in the cortex, better than null models that assumed thalamocortical interactions do not carry task-specific information. Furthermore, past studies that adopted similar activity flow mapping approaches have shown that including all regions across the whole brain can predict task-specific activity patterns with very high accuracy (Cole et al., 2016). Our study tested a different question to evaluate the predictive power of the thalamus relative to other brain regions. Critically, we found that the thalamocortical model outperformed comparison models constructed with other cortical and subcortical structures, including frontal (the insula and the inferior frontal sulcus), parietal, and striatal regions previously implicated in adaptive control of tasks (Badre and Nee, 2018; Ito et al., 2022). These results suggest that thalamocortical interactions likely exert a strong influence on cortical activity that may instantiate cognitive representations.

Placing the thalamus in a central position for influencing cortex-wide cognitive activity has several benefits. For instance, the thalamus consists of two classes of thalamocortical projection cells, “matrix” and “core” cells, where core cells target specific granular cortical structures, and matrix cells diffusely project to multiple brain regions (Jones, 2001). Given that different brain regions may be encoding different perceptual and cognitive information (Christophel et al., 2017), selectively modulating thalamic activity may facilitate targeted activations of cortical representations via associated thalamocortical interactions. The diffuse projection pattern of matrix cells in the thalamus may simultaneously activate multiple cortical regions and promote inter-system integration when required (Hwang and D’Espostio, 2022; Jones 2001). Furthermore, the thalamus is modulated by inputs from other subcortical structures, such as the reticular nucleus, the basal ganglia, the superior colliculus, and the cerebellum (Bostan and Strick, 2018; Shine, 2020). This anatomical feature suggests that the thalamus may be an efficient target for exerting neuromodulatory (e.g., dopaminergic) influences on cognitive processes when required (Castro-Alamancos and Gulati, 2014; Garrett et al., 2022). Several computation models have leveraged these characteristics to explain the functions of basal-ganglia-thalamic circuits for higher-order cognitive functions (Jaramillo et al., 2019; O’Reilly and Frank, 2006).

Another important observation was that the anterior, medial, and posterior thalamus exhibited “task hub” property, that is, are more flexibly recruited by multiple thalamic activity components. These subregions overlapped with thalamic FC hubs that we previously identified (Hwang et al., 2017). In the current study, we found that thalamic task hubs were most strongly coupled with the frontal and parietal associative cortices (Figure 2B). Intriguingly, these frontoparietal regions have been hypothesized to be part of a flexible multiple-demand system (Cole et al., 2013; Duncan, 2010; Ito et al., 2022), and thus, these thalamic subregions may be part of a core system whose functions are required to perform specific computations that are common across many different task contexts. One strong test of the behavioral relevance of these thalamic task hubs is to examine the effects of lesions. If thalamic task hubs are part of a domain-general core, lesioning task hubs should be associated with broad behavioral deficits not confined to a specific behavioral domain. Indeed, we found that simulated lesions to thalamic task hubs impaired prediction accuracy in cortical activity pattern across tasks, and stroke lesions to the anterior-medial thalamus in human patients were associated with behavioral deficits across functional domains (Hwang et al., 2021, 2020a). At this point we are agnostic to the specific function these thalamic task hubs perform. One hypothesis, suggested by findings from rodent models, is that the medial thalamus excites task-relevant and inhibits task-irrelevant cortical activity that encode working memory representations (Mukherjee et al., 2021; Rikhye et al., 2018). Thus, in human subjects, thalamocortical interactions between thalamic task hubs and frontoparietal systems may control task representations that bind sensory, motor, and contextual information that are necessary for adaptive behavior across tasks (Schumacher and Hazeltine, 2016). This is an open question that should be evaluated in future studies.

On important limitation of our study is that our thalamocortical activity flow model cannot establish the directionality of thalamic activity influencing cortical activity. It is more likely that brain systems instantiate cognitive representations via recurrent interactions among cortical regions and the thalamus, as well as integrating inputs from other brain structures. As discussed above, including all brain regions into the activity flow model can substantially improve model performance in predicting task-specific activity patterns (Cole et al., 2016). Nevertheless, our findings suggest that, across the whole brain, the thalamus likely has one of the strongest contributions to cortical task activity. Another important limitation is that the posterior thalamus exhibited strong task hub property in the MDTB dataset but not the N&N dataset. We suspect this could be related to differences in behavioral tasks across the two datasets, and speculatively, to the increased noise in the N&N dataset, given the lower number of observations per task and lower subcortical signal-to-noise ratio. One strong test of mapping task hubs is the lesion methods, and in our previous study (Hwang et al., 2021) we did not have good lesion coverage on the pulvinar nucleus. Future studies should test whether patients with pulvinar lesions have cognitive deficits beyond visual attention tasks (Snow et al., 2009).

To conclude, our study reported two important organizational principles of how thalamic task-evoked activity and thalamocortical interactions support cognitive tasks. Thalamic task-evoked activity has a low-dimensional structure, and thalamocortical interactions have a strong influence on generating task-specific cortical activity patterns. These results highlight how the thalamus is in a central position for supporting cognitive task representations.

## Methods

### Datasets

We analyzed two datasets, the multi-domain task battery dataset (MDTB dataset [King et al., 2019] at https://openneuro.org/datasets/ds002105/) and the Nakai and Nishimoto dataset (N&N dataset [Nakai and Nishimoto, 2020] at https://openneuro.org/datasets/ds002306/). We selected these datasets because both include fMRI data from subjects that performed a large number of tasks across functional domains (Figure 1A). The MDTB dataset included fMRI data from 21 subjects (13 women, 8 men; mean age = 23.28 years, SD = 2.13; we excluded 3 out of the original 24 subjects that did not have all tasks available after removing high noise datapoints). MRI Data were collected on a 3T Siemens Prisma with the following parameters: repetition time (TR) = 1 s; 48 slices with 3 mm thickness; in-plane resolution 2.5 × 2.5 mm^2^; multi-band acceleration factor = 3; in-plane acceleration factor =2; echo time and flip angle were not reported in the original paper. Structural T1 images were acquired using magnetization-prepared rapid acquisition gradient echo sequence (MPRAGE; field-of-view = 15.6 × 24 × 24 cm^3^, at 1 × 1 × 1 mm^3^ voxel size). The N&N dataset included fMRI data from 6 subjects (2 females, 4 males, age range 22-33 years) collected on a 3T Siemens Tim Trio system. Functional MRI data were collected with the following parameters: TR = 2 s; 72 slices with 2mm slice thickness; in-plane resolution = 2 × 2 mm2; echo time = 30 ms; flip angle = 62 degrees; multiband factor = 3. Structural MRI data were collected with a MPRAGE sequence (TR = 2530 ms; TE = 3.26 ms; flip angle=9 degrees; FOV=256 × 256 mm^2^; voxel size=1×1×1mm^3^).

### MRI data preprocessing

Both datasets were preprocessed using fMRIPrep version 20.1.1 (Esteban et al., 2019) to reduce noise and transform data from subject native space to the ICBM 152 Nonlinear Asymmetrical template version 2009c for group analysis (Fonov et al., 2011). Preprocessing steps include bias field correction, skull-striping, co-registration between functional and structural images, tissue segmentation, motion correction, and spatial normalization to the standard space. As part of preprocessing, we obtained nuisance regressors including rigid-body motion estimates, cerebral spinal fluid (CSF), and white matter (WM) noise components from the component-based noise correction procedure (Behzadi et al., 2007). These nuisance regressors were entered in the regression model to reduce the influences from noise and artifacts (see below). We did not perform any spatial smoothing.

### Task-evoked responses

The MDTB dataset contained 24 cognitive tasks and a resting condition collected across four scanning sessions; each session consisted of 8 runs. Each task started with a 5 second instruction and was followed by 30 seconds of continuous task performance. The N&N dataset contained 103 tasks collected across 18 MRI scanning runs in 3 days. Each task trial lasted 6 to 12 seconds where task instruction was on screen throughout the trial, and each task repeated 12 times distributed across runs. For a complete list of behavioral tasks, see Figure 1A and Figure 1-Supplementary File 1. The detailed design of each task for both datasets were reported in King et al., 2019, and in Nankai & Nishimoto, 2020.

We employed a voxel-wise general linear modeling (GLM) approach to estimate task-evoked responses, using AFNI’s 3dDeconvolve (Cox, 1996). For every voxel, a generalized least squares regression model was constructed and fitted to the preprocessed timeseries with the following regressors: task regressors, rigid body motion regressors and their derivatives, and the top 5 CSF and WM noise components. High motion volumes (framewise displacement > 0.2 mm) were removed from data analyses via the censoring option, and subjects with tasks with more than 40% of the data censored were dropped from further analyses (3 MDTB subjects dropped). For each subject, all imaging runs were concatenated, and signal drifts were modeled separately for each run, including run-specific constant and polynomial regressors. All task regressors were created by convolving a gamma hemodynamic response function with the stimulus duration. The stimulus duration was determined by the specific block or trial design sequence of each task (See Figure 1-Supplementary File 1 for the specific duration used for all tasks). For the MDTB dataset, several tasks contained different sub-conditions, and the GLM estimates of sub-conditions were averaged to obtain one estimate per task. Repeating our analyses without averaging sub-conditions did not change our results. For the MDTB dataset, the 5 s instruction period at the start of each task block was modeled but not included into subsequent analyses. For the N&N dataset, the task instruction was presented simultaneously during all trials, and as a result it could not be separated from the GLM analyses.

### Low-dimensional organization of task-evoked activity in the thalamus

To probe the low-dimensional organization of task-evoked activity in the thalamus, we first applied a thalamus mask from the Harvard-Oxford subcortical atlas, which mask was used to extract task-specific evoked response estimates from 2445 thalamus voxels. These estimates were then compiled into a task-by-voxel evoked activity matrix for each subject. This evoked activity matrix was then z-scored by subtracting the grand mean from the whole matrix and divided by the standard deviation across all elements. A principal component analysis decomposed the evoked activity matrix into a linear summation of a voxel-by-component weight matrix multiplied by a component-by-task loading matrix (Figure 1A). The voxel-by-component weight matrix can be conceptualized as sets of basis patterns of thalamic activity components engaged by different tasks. The loading matrix described the relationship between each voxel-wise component map and tasks. We then summated the percentage of variances explained to determine how many components were required to explain more than 50% of the variance observed in the evoked activity matrix. We performed the PCA analysis with and without averaging the evoked activity matrix across subjects. We averaged the matrix before PCA to visualize the voxel patterns of each component and repeated the PCA analysis without averaging separately for each subject to test whether the low-dimensional structure can be replicated at the level of individual subjects.

### Thalamic task hubs

Some thalamic subregions could participate in multiple tasks across functional domains, exhibiting “task hub” properties. To map these task hub regions, for every voxel we calculated a *WxV* metric:

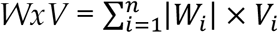

Where |*W*_*i*_| is the absolute value of weight for component *i, V*_*i*_ is the percentage of variance explained for component *i*, and n is the number of components. We reasoned that a task hub region would be more strongly recruited by components that explained a large amount of the variances in the evoked activity matrix, exhibiting a higher *WxV* estimate.

To compare the task hubs to the FC hubs that we had mapped previously (Hwang et al., 2017), we first obtained a thalamocortical FC matrix using principle component linear regression (Ito et al., 2017) to estimate patterns of FC between each thalamic voxel and 400 cortical ROIs (Schaefer et al., 2018). We estimated this FC matrix using residuals after task regression. Note that one advantage of principle component linear regression is that its estimates are similar to partial correlation, accounting for shared variances among signals. We then calculated a connector hub metric participation coefficient for each voxel (Gratton et al., 2012):

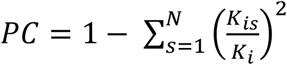

where *K*_*i*_ is the sum of total FC weight for voxel *i, K*_*is*_ is the sum of functional connectivity weight between voxel *i* and the cortical network *s*, and *N* is the total number of networks. To perform this calculation, we assigned the 400 cortical ROIs to seven cortical functional networks, i.e., to the FP, DF, CO, DA, limbic, SM, and visual networks (Schaefer et al., 2018; Yeo et al., 2011). We calculated participation coefficient values across a range of density thresholds of the thalamocortical FC matrix (density = 0.01–0.15) and averaged across thresholds.

To determine which cortical regions showed the strongest coupling with thalamic task hubs, we calculated the dot product between *WxV* and thalamocortical FC matrix, yielding a project task hub matrix on the cortical space:

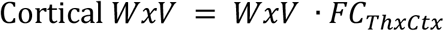

where *WxV* is the 2445 voxel-wise task hub metrics, and *FC*_*ThxCtx*_ is the 2445 × 400 thalamocortical FC matrix.

### Thalamocortical activity flow mapping

To determine whether thalamocortical interactions can transform thalamic task-evoked activity to cortical task activity, we modified the activity flow mapping procedure (Cole et al., 2016; Ito et al., 2017), and for each task and each subject we calculated:

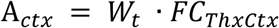

where *W*_*t*_ is the evoked response estimate for every thalamic voxel, *FC*_*ThxCtx*_ is the thalamocortical FC matrix, and A_*ctx*_ is the predicted cortical activity pattern. We repeated this analysis for every task, and the predicted cortex-wide activity patterns were then empirically compared to the observed activity patterns using Pearson correlation. In addition to comparing individual task activity patterns, we also compared the predicted versus observed representational geometry. Representation geometry seeks to capture the topography of task representations by characterizing the multivariate distance of activity patterns among tasks. We calculated the representational dissimilarity matrix using Spearman correlation. Predicted and observed representational dissimilarity matrices were compared using Kandal’s Tau (Nili et al., 2014).

To evaluate our thalamocortical activity flow mapping procedure, we compared its results to two different null models and to comparison models. The first null model randomly shuffled the voxel order in *W*_*t*_, which assumed no task-specific information in thalamic evoked activity pattern. The second null model set *W*_*t*_ to a uniform vector, which assumed that A_*ctx*_ was entirely determined by the pattern in *FC*_*ThxCtx*_. We constructed a series of comparison models by repeating the same analysis on other source ROIs, including the hippocampus, the caudate, the putamen, and the pallidus, and 100 cortical ROIs (Schaefer et al., 2018) that were similar in size to the thalamus. For these comparison models, we replaced *W*_*t*_ with voxel-wise evoked responses from other source ROIs, and recalculated the FC matrix by calculating FC between every voxel in the source ROI and 400 cortical ROIs using PCA linear regression.

### Lesion analyses

We performed additional simulated and empirical lesion analyses to test which thalamic subregions contribute to predicting cortical task activity. For simulated lesions, we ranked thalamic voxels by their task hub (*WxV*) metrics, then set the evoked responses in 20% of thalamic voxels to zero and repeated the thalamocortical activity flow analysis. We repeated this virtual lesioning procedure for voxels ranking from 0% to 80% in steps of 1%, window size of 20%, and calculated the effects of virtual lesioning of thalamic voxels on predicting both individual task activity pattern and representational geometry. The percentage of prediction reduction (relative to the original observed value) after virtual lesioning was assigned to each voxel to construct voxel-wise maps that depict the effects of simulated lesions.

We then compared the effects of simulated lesions to lesions observed in human patients with focal thalamic lesions. Details of these patients were reported in our previous study (Hwang et al., 2021). Briefly, these patients were selected from the Iowa Neurological Patient Registry, and had focal lesions caused by ischemic or hemorrhagic strokes restricted to the thalamus (age = 18-70 years, mean = 55.8 years, SD = 13.94 years, 13 males). The lesion sites of these patients were manually traced and normalized to the MNI-152 template using a high-deformation, non-linear, enantiomorphic, registration procedure that we described in detail in our previous papers (Hwang et al., 2021, 2020a).

We further tested whether voxels associated with stronger virtual lesioning effects overlapped with lesion sites associated with more pronounced behavioral deficits in human stroke patients. Behavioral deficits were assessed using a set of standardized neuropsychological tests: (1) executive function using the Trail Making Test Part B (TMT Part B); (2) verbal naming using the Boston Naming Test (BNT); (3) verbal fluency using the Controlled Oral Word Association Test (COWA); (4) immediate learning using the first trial test score from the Rey Auditory-Verbal Learning Test (RAVLT); (5) total learning by summing scores from RAVLT, trials one through five; (6) long-term memory recall using the RAVLT 30-minute delayed recall score; (7) long-term memory recognition using the RAVLT 30-minute delayed recognition score; (8) visuospatial memory using the Rey Complex Figure delayed recall score; (9) psychomotor function using the Trail Making Test Part A (TMT Part A); and (10) construction using the Rey Complex Figure copy test. To account for age-related effects, all test scores were converted to age-adjusted z-scores using the mean and standard deviation from published population normative data. We determined the functional domain that each test assessed, described in Neuropsychological Assessment (Lezak et al., 2012). Twelve out of 20 patients had significant impairment (z < -1.645) reported in more than two functional domains, and thus were classified as the multiple-domain (MM) patient group. The rest of the 8 patients had significant impairment in one or fewer domains, and classified as the single-domain (SM) group. We predicted that lesion sites in the MM group would show stronger virtual lesioning effects, when compared to lesion sites in the SM group.

## Acknowledgments

E.S. and K.H. were supported by National Institute of Mental Health R01MH122613. The content is solely the responsibility of the authors and does not represent the official views of the National Institutes of Health.

## Code and data availability statement

Code and data are available at https://github.com/HwangLabNeuroCogDynamics/ ThalamicTaskHubs

---

**Figure 1 – Figure Supplement 1.**
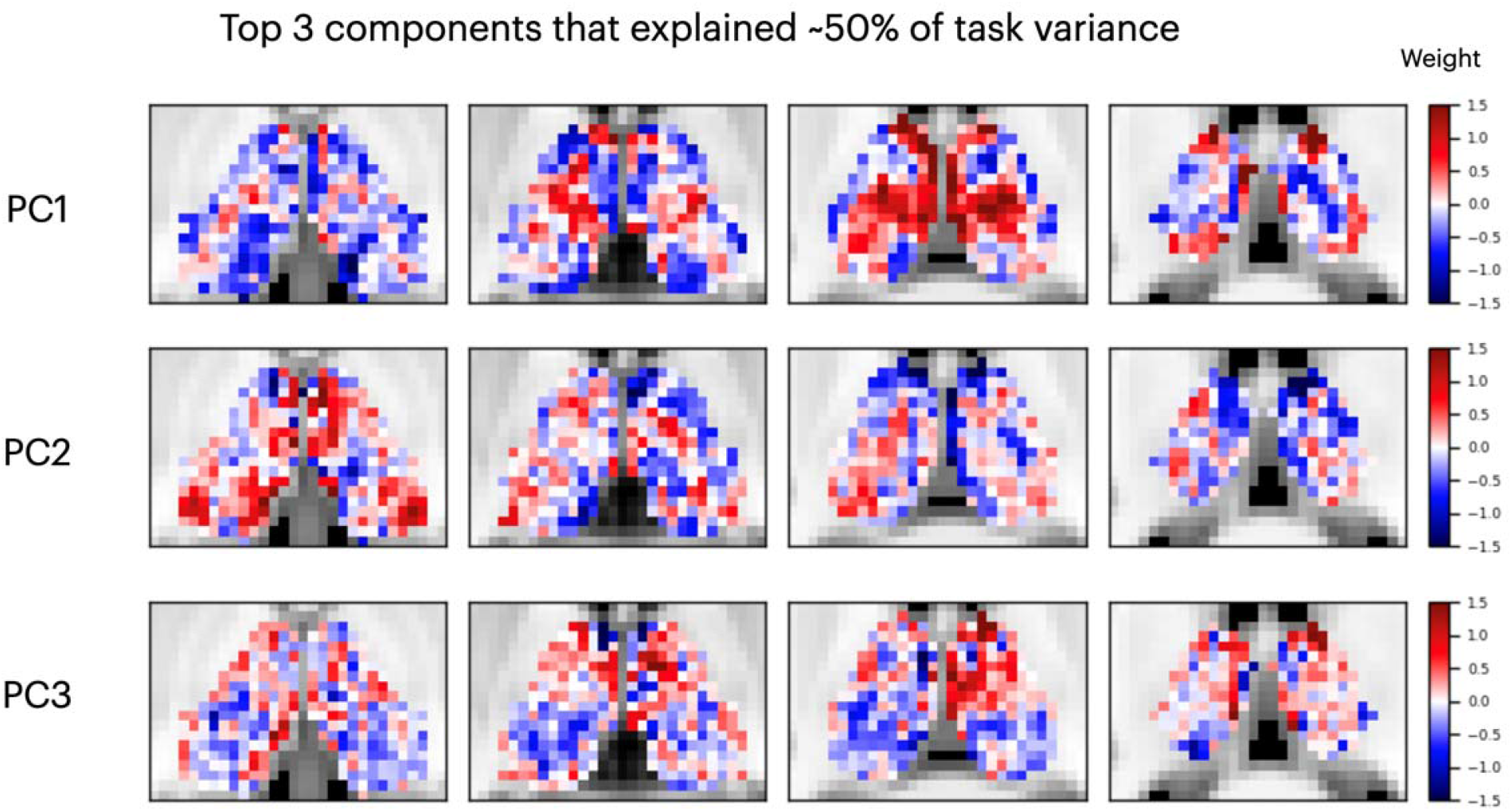
Spatial topography of the top three components from the N&N dataset that explained about 50% of the variance in the group averaged task activity matrix.

**Figure 1 – Figure Supplement 2.**
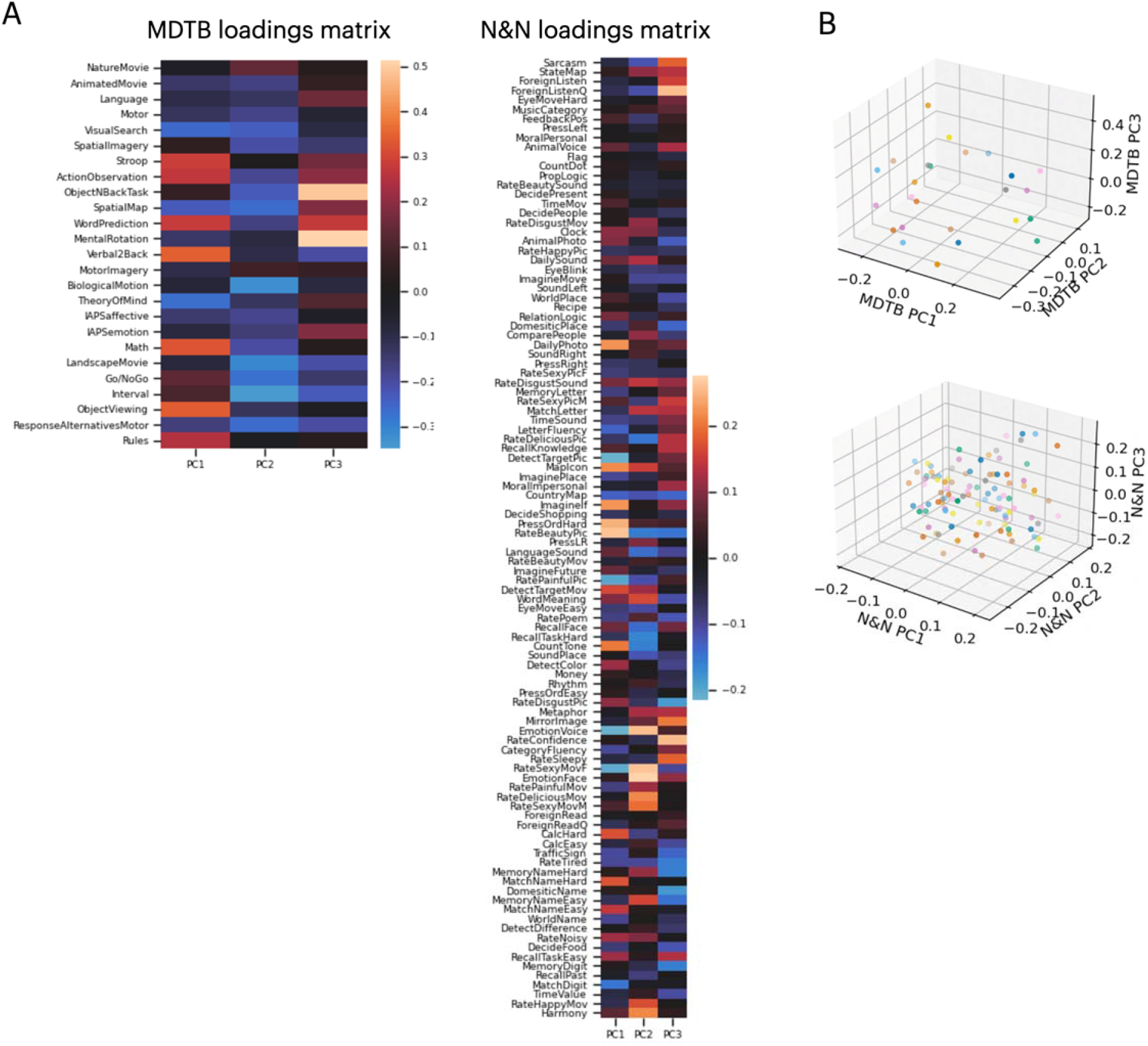
Loadings between individual tasks and thalamic activity components. (A) Loadings matrix for the top 3 activity components and individual tasks for both datasets. (B) Low-dimensional embedding of task-evoked responses, plotted by task loadings on each activity component. Each colored dot represents a single task.

## Notes

### Competing Interest Statement

The authors have declared no competing interest.

